# A novel, anatomy-similar in vitro model of 3D airway epithelial for anti-coronavirus drug discovery

**DOI:** 10.1101/2021.03.03.433824

**Authors:** Yaling Zhang, Dingailu Ma, David Jitao Zhang, Xin’an Liu, Qihui Zhu, Li Wang, Lu Gao

## Abstract

SARS-CoV-2 and its induced COVID-19 remains as a global health calamity. Severe symptoms and high mortality, caused by cytokine storm and acute respiratory distress syndrome in the lower respiratory airway, are always associated with elderly individuals and those with comorbidities; whereas mild or moderate COVID-19 patients have limited upper respiratory flu-like symptoms. There is an urgent need to investigate SARS-CoV-2 and other coronaviruses replication and immune responses in human respiratory systems. The human reconstituted airway epithelial air-liquid interface (ALI) models are the most physiologically relevant model for the investigation of coronavirus infection and virus-triggered innate immune signatures. We established ALI models representing both the upper and the lower respiratory airway to characterize the coronavirus infection kinetics, tissue pathophysiology, and innate immune signatures from upper and lower respiratory tract perspective. Our data suggested these in vitro ALI models maintain high physiological relevance with human airway tissues. The coronavirus induced immune response observed in these upper and lower respiratory airway models are similar to what has been reported in COVID-19 patients. The antiviral efficacy results of a few promising anti-coronavirus drugs in these models were consistent with previous reports and could be valuable for the human dose prediction. Taken together, our study demonstrates the importance of 3D airway epithelial ALI model for the understanding of coronavirus pathogenesis and the discovery and development of anti-coronavirus drugs.

## Introduction

Global pandemic of coronavirus disease 2019 (COVID-19) caused by SARS-CoV-2 infection is a serious global health threat. As of one year after its outbreak in Wuhan, Hubei Province, China (WHO COVID-19 weekly epidemiological update), more than 100 million cases and nearly 2.3 million deaths worldwide have been confirmed. Most patients with COVID-19 exhibit mild to moderate symptoms of upper respiratory tract. However, nearly 15% of patients progress to severe lower respiratory tract symptoms such as pneumonia and about 5% eventually develop lung acute respiratory distress syndrome (ARDS) or cytokine storm syndrome (Huang et al., 2020; Xu et al., 2020b). The odds of severe symptoms are higher with elder people (Meftahi et al., 2020). The fatality rate for COVID-19 patients aged ≥ 60 years was 63.6% from the initial data of Wuhan, which is significantly higher than that of patients < 60 years (Chen et al., 2020; Liu et al., 2020).

The human respiratory tract is a highly specialized organ comprising upper and lower tracts, both of which are susceptible to SARS-CoV-2 infection (Lui et al., 2020). A deep understanding of SARS-CoV-2 pathogenesis and its induced clinical features require comprehensive characterization of the virus infection and the associated host response. However, Vero E6 cell-based infection assay, the most frequently used system for the assessment of anti-SARS-CoV-2 inhibitors, has limitations in physiological relevance. Several reported in vitro lung models can complement animal models for the deep understanding of pathological mechanisms of respiratory viruses (Miller and Spence, 2017; Takayama, 2020), among which the human reconstituted airway epithelial air-liquid interface (ALI) models are highly relevant and useful tools for pre-clinical studies of COVID-19 (Mulay et al., 2020). Human bronchial epithelial cells (NHBE) or small airway epithelial cells (SAEC) differentiated in the air/liquid interphase represents the human upper or lower respiratory tract, respectively. These ALI models reflect similar human respiratory epithelium anatomy features, including the main structure, function and innate immune responses in the early stages of virus infection and constitute robust surrogates to study respiratory viral infections (Boda et al., 2018). A few studies characterized the replication of SARS-CoV-2 in the proximal airway NHBE ALI (Hao et al., 2020; Zhu et al., 2020). In contrast, the investigation of SARS-CoV-2 replication in the human small airways has not been reported. In line with the severe symptoms of COVID-19 in lower respiratory tract, the detrimental outcomes of virus infection often related with immune dysregulation in lower respiratory tract, termed as “cytokine storm” (Song et al., 2020). Accumulating evidence suggests that a subgroup of patients with severe COVID-19 might have a cytokine storm syndrome (Huang et al., 2020; Pedersen and Ho, 2020; Zhou et al., 2020). Therefore, a thorough characterization of both the upper and lower respiratory airway in the context of coronavirus infection is in urgent need for the understanding of COVID-19 immunopathology and anti-coronavirus drug discovery.

Here, we reconstituted the human upper (NHBE) and lower (SAEC) respiratory airway from primary epithelial cells of bronchial and small airway origin and characterized their structural, functional and innate immune features. We also investigated coronavirus infection kinetics, tissue pathology, and transcriptional immune signatures induced by the coronavirus infection in these ALI models. Our data suggest that SAEC ALI, a novel 3D airway epithelial ALI model, represents comparable anatomy features and gene expression profile of human airway tissues. Besides, innate immune responses and cytokine profile generated from NHBE and SAEC models are similar to the data from patients in both upper and lower respiratory tracts. We further validated the potential of SAEC ALI for evaluating antiviral drugs against SRAS-CoV-2 and other coronaviruses. Our results provide important insights into coronaviruses kinetics, pathogenesis, and innate immune responses, and suggest the combination treatment of the polymerase inhibitor and the 3C-like protease inhibitor would be potential beneficial for the COVID-19.

## Results

### Characterization of the in vitro 3D airway ALI model of differentiated human bronchial and small airway epithelial cells

Recent studies focused on SARS-CoV-2 pathogenesis investigation or antiviral drug evaluation were mainly rely on cell lines or primary epithelial cells (Choy et al., 2020). However, these monolayer cultures consist only basal cells and could not fully sustain viral replication and innate immune activation. Therefore, two ALI models, NHBE ALI representing part of the upper respiratory tract and SAEC ALI representing part of the lower respiratory tract, were established to investigate the coronavirus replication kinetics and innate immune activation in human airways A fully differentiated pseudostratified epithelial layer was determined to be achieved by 4 weeks post air-lifted (Fig. 1A and 1B). The ALI culture thickness of both SAEC and NHBE at 4 weeks were around ten times thinner to what was reported for in vivo human airway epithelium, and NHBE ALI was nearly three times thicker than SAEC ALI culture (Fig. 1A and 1B) (Weibel, 1990). The differentiation capability of the primary airway epithelial cells was largely associated with the age of donor. The younger the donor is, the higher the potential to be differentiated (Fig. 1B). The cilia beating of SAEC ALI and NHBE ALI indicates the successful differentiation of the ciliated epithelium (Fig. S1A,). The in vitro airway ALI model has tight junctions and distinct apical and basolateral membranes. High transendothelial electrical resistance (TEER) value implies the formation of an impermeable tight junction at the apical side. The TEER value of SAEC and NHBE ALI was measured every 2-3 days for up to 60 days. The TEER value for NHBE ALI reached to a maximum value of about 1400 Ω*cm^2^ at 2-3 weeks post air-lifted and gradually dropped to around 300 Ω*cm^2^ after 35 days post air-lifted. In contrast, the TEER value of SAEC ALI did not decrease after reaching plateau, and the plateau value is ~300 Ω*cm^2^, which is 3.6-fold less compared with NHBE ALI (Fig. 1C). The data demonstrate the upper and lower respiratory airways differ in morphology and differentiation, which might lead to differential outcomes upon viral infections.

**Figure 1.**
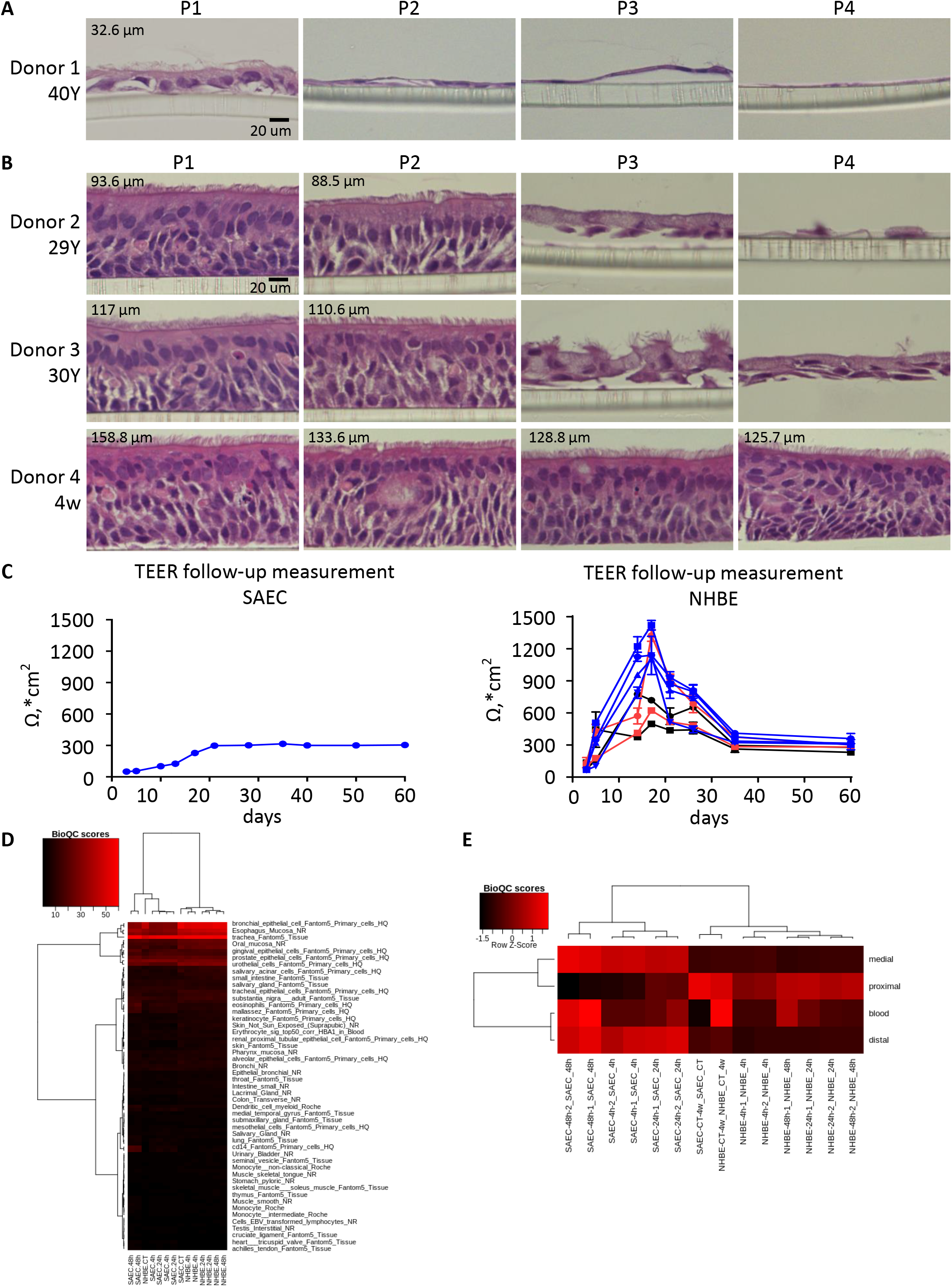
Characterization of in vitro 3D airway ALI model of differentiated human bronchial and small airway epithelial cells. **A**, Human primary bronchial airway epithelial cells originated from different donors (29 years old; 30 years old and 4 weeks fetal) and different passages (P1-P4) were differentiated at air-liquid interface. H&E staining of the 3D cell culture was performed at 5 weeks post air-lift. The thickness of the culture was measured and labeled at the upper-left corner of the image. **B**, Human primary small airway epithelial cells originated from donor age at 40 years old at passage 1 was differentiated at air-liquid interface. H&E staining of the 3D cell culture was performed at 5 weeks post air-lift. The thickness of the culture was measured and labeled at the upper-left corner of the image. The shown photos are representative of 3 independent experiments. Scale bar denotes 20 μm. **C**, Resistance values (Ω) was measured at the indicated days and converted to TEER (Ω*cm2) value. Symbols shown in the NHBE ALI culture: Sciencell P1 (blue circles); Sciencell P2 (blue squares); Sciencell P3 (blue triangles); Sciencell P4 (blue inverted triangles); Lonza P1 (red circles); Lonza P2 (red squares); ATCC P1 (black circles); ATCC P2 (Black aquares). Symbols shown in the SAEC ALI culture: Lonza P1 (blue circles). Data are representative of 3 independent experiments. **D**, Heatmaps depict the bioQC scores derived from SAEC and NHBE ALI models for testing for the enrichment of signatures in different tissues. BioQC scores are defined as log10-transformed one-sided p-values of Wilcoxon-Mann-Whitney test. Red and black indicate high and low scores respectively. **E**, Standardized BioQC scores of SAEC and NHBE ALI models before and after virus infection are visualized in heatmaps. The enrichment analysis was performed using signature genes derived from different regions of lung tissues. Red and black indicate high and low scores respectively. Raw scores above the mean have positive standard scores, while those below the mean have negative standard scores.

Next, the transcriptome profile of SAEC and NHBE ALI cultures was examined by RNA sequencing experiments. To characterize the similarity between the in vitro ALI model and the in vivo human respiratory airway tissues, we analyzed the enrichment of the signature genes derived from lung scRNA-seq data (Deprez et al., 2020; Travaglini et al., 2020), and found that SAEC and NHBE ALI samples were indeed enriched with the signature genes in human lung epithelial cells (Fig. 1D and Fig. S2A). Although we could hardly distinguish SAEC from NHBE based on the signature genes in single/specific cell subtype (Fig. S2B), we could clearly observe that SAEC samples were significantly closer to distal airways, while NHBE samples were nearer to proximal airway cells when grouping these cells according to the locations (Fig. 1E). In summary, our data demonstrate the successful reconstitution of the in vitro 3D airway ALI model with both upper and lower respiratory airway epithelial cells.

### The replication of coronaviruses in human upper and lower respiratory airways

We next characterized the replicative capacities of three human coronaviruses SARS-CoV-2, HCoV-OC43 and HCoV-229E in human upper and lower respiratory airway ALI models. SAEC and NHBE ALIs were infected with SARS-CoV-2 or HCoV-OC43 at a multiplicity of infection (MOI) of 0.1, and infected with HCoV-229E at MOI of 0.0001 to 0.01 from the apical side and the viral yield at the same side was quantified at different time points post-infection. SARS-CoV-2 displayed a time-dependent increase in viral titers and the peak titer was observed at 72 hours post infection (hpi) in both NHBE and SAEC ALI. (Fig. 2A). The growth kinetics of SARS-CoV-2 in these two models was indistinguishable. OC43 showed a similar growth curve as SARS-CoV-2 in SAEC ALI. However, in NHBE ALI, no increase in viral yield was observed (Fig. 2B), suggesting the replication of OC43 in NHBE ALI might be abortive. Compared with SARS-CoV-2 and HCoV-OC43, 10 to 1000-fold less inoculum of HCoV-229E was used because even at the MOI as low as 0.0001, severe cytopathic effect was observed for SAEC ALI within 48 hrs. At all tested MOIs, HCoV-229E viral production at the epithelial apical surface of SAEC ALI increased sharply upon virus addition, reached the plateau at 48 hpi, and then declined rapidly (Fig. 2C left panel, Fig. S3A), suggesting the replication of HCoV-229E in SAEC ALI. Continuous increase of the virus titer was observed for all MOIs in NHBE ALI from day 1 to day 5 post infection (Fig. 2C right panel, Fig. S3B), with a peak viral titer similar to that in SAEC ALI.

**Figure 2.**
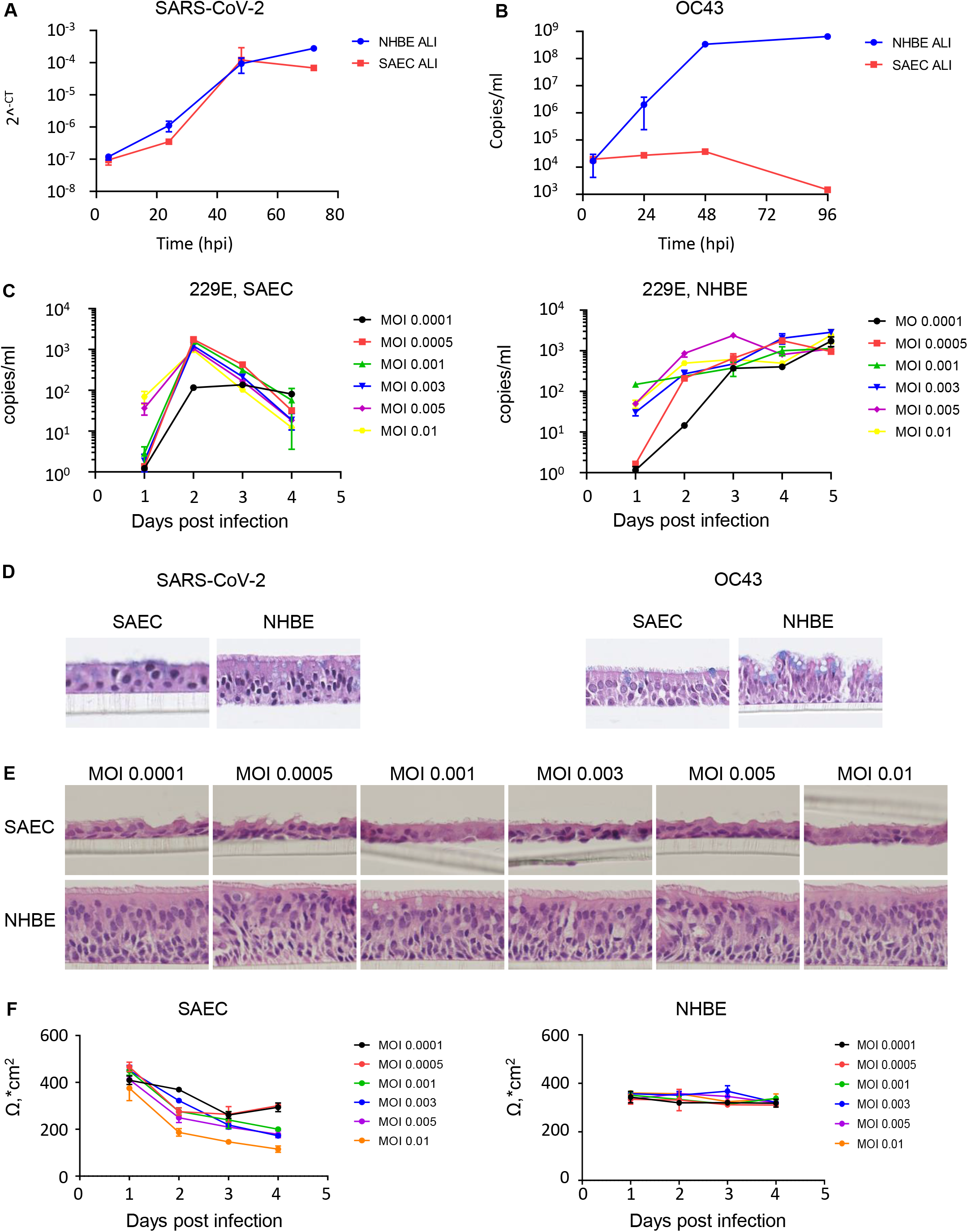
Different replication kinetics and tissue pathophysiology of coronavirus in human upper and lower respiratory airway. **A**, SARS-CoV-2 replication kinetics measured by qPCR at indicated time points in NHBE ALI (blue circle) and SAEC ALI (red square) culture. **B**, OC43 replication kinetics measured by qPCR at indicated time points in NHBE ALI (blue circle) and SAEC ALI (red square) culture. **C**, 229E replication kinetics measured by qPCR with different MOIs at indicated time points in SAEC ALI and NHBE ALI culture. **D**, H&E staining of the 3D cell culture at the end of infection of SARS-CoV-2 (72 hpi) or OC43 (96 hpi). **E**, H&E staining of the 3D cell culture at the end of 229E infection with different MOIs in SAEC and NHBE ALI cultures. **F**, Trans-epithelial resistance (TEER in Ω, *cm^2^) of 229E-infected SAEC or NHBE was measured at each time point. All data are presented as mean ± SD of 3 independent experiments.

At the end of the experiment, ALI cultures were processed for H&E staining to understand the morphology remodeling caused by these viral infections. The SARS-CoV-2 infected SAEC ALI had dramatically less ciliated cells, reduced thickness (Fig. 2D, left panel), and non-observable cilia beating (data not shown). Interestingly, no significant morphology changing was observed in NHBE ALI after SARS-CoV-2 infection (Fig. 2D, left panel), suggesting the host response in NHBE ALI might differ from that in SAEC ALI. HCoV-229E showed very similar pathogenesis as SARS-CoV-2 (Fig. 2E). Coincident with the rapid decline in viral titer, the HCoV-229E-infected SAEC ALI had significantly less ciliated cells, reduced thickness, and a reduction in TEER (Fig. 2E-F, Fig. S1B, S3C), and no obvious characteristic cytopathic effect was found in NHBE ALI (Fig. 2E-F, Fig. S3C). The phenotypic differences in NHBE and SAEC ALI epithelia pathology was not elicited by the viral load difference, because the apical surface virus titer in NHBE ALI was even higher than that in SAEC ALI (Fig. 2C). OC43 infection resulted in an opposite phenomenon that the epithelial integrity of NHBEALI was damaged, but SAEC ALI was intact (Fig. 2D, right panel), which is well in line with its viral replication kinetics in these two models (Fig. 2B).

Taken together, these data demonstrate SARS-CoV-2 and HCoV-229E could efficiently replicate in both NHBE and SAEC ALI and produce infectious virions, and HCoV-229E shares high degree of pathogenesis similarity with SARS-CoV-2 in both upper and lower respiratory tract of human airway.

### Coronavirus infection elicits differential cytokine profiles in upper and lower respiratory airway

Next, we evaluated the production of a panel of 33 cytokines in both NHBE and SAEC ALI infected by HCoV-229E for 24 hours to further understand the pathophysiological differences between upper and lower respiratory airways. Upon HCoV-229E infection, 18 cytokines were regulated, among which the number of cytokines that are either upregulated or downregulated from the apical side were remarkably less than the basal compartment in both NHBE and SAEC ALI (Table 1), however most of the pro-inflammatory cytokines were up-regulated at apical whereas down-regulated in the basal compartment (Table 1). These observations indicate the cytokine release in respiratory tracts is polarized and the level of the proinflammatory cytokines in the pulmonary side is much higher than the blood circulation at early time of infection.

**Table 1.**
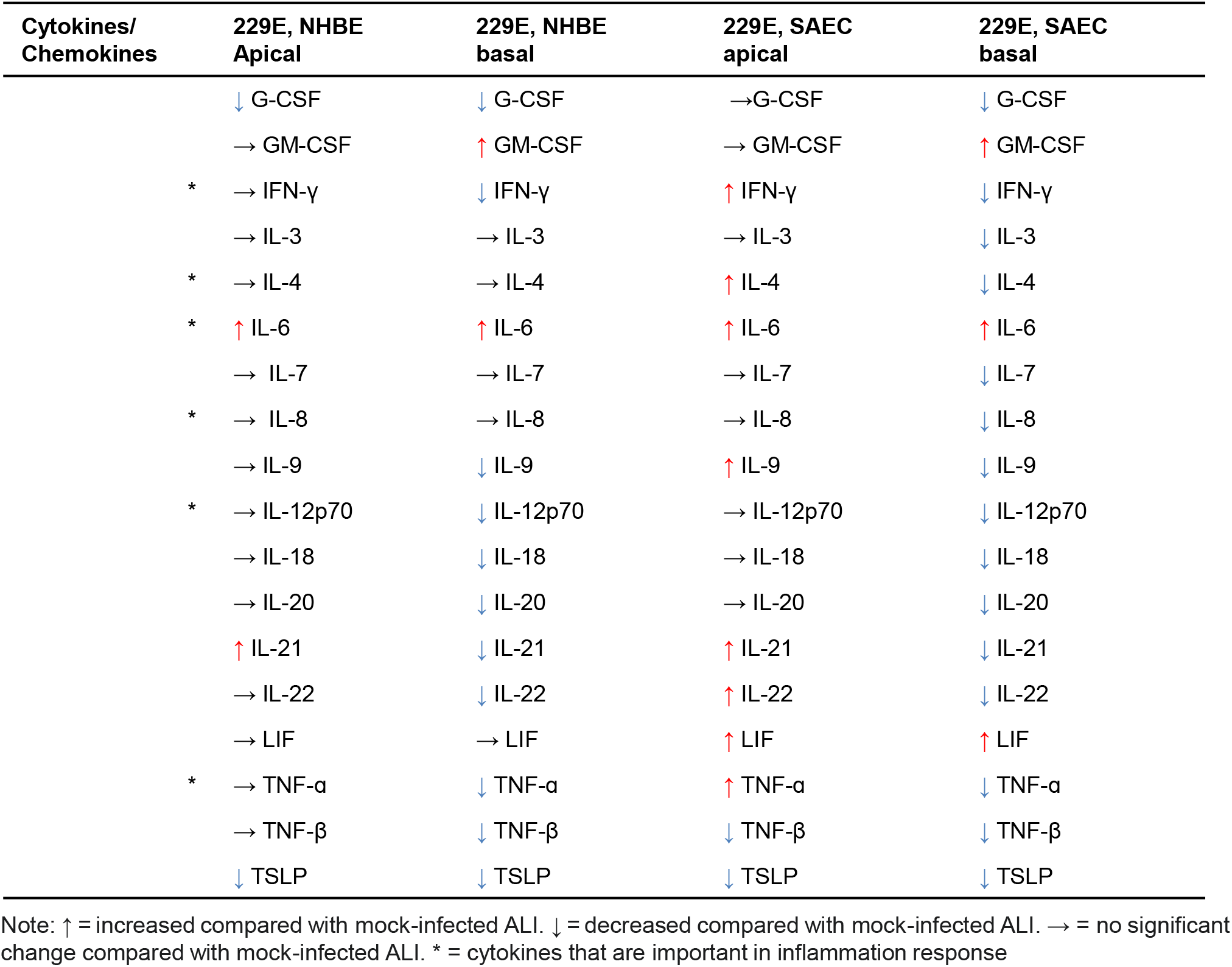
Cytokines & chemokine production profiles of HCoV-229E-infected NHBE and SAEC ALI

When all the upregulated cytokines from the apical side of NHBE and SAEC ALI were analyzed side by side, we observed that a total of 8 cytokines (IFN-γ, IL-4, IL-6, IL-9, IL21, IL22, LIF and TNF-α). were upregulated in the apical side of SAEC ALI after HCoV-229E infection in a MOI-dependent manner (Fig. 3A), however only 2 cytokines (IL-6 and IL21) were upregulated in NHBE ALI (Fig. 3B). These results are consistent with cytokine analysis data from COVID-19 patients (Huang et al., 2020), and demonstrate the HCoV-229E infection triggered an aggressive induction of cellular immune response in SAEC but not in NHBE ALI, which is correlated with the observation of the differential tissue damage caused by HCoV-229E infection in NHBE and SAEC ALI. IL-6 appeared to be highly interesting as it was well known to play a critical role in COVID-19 disease exacerbation. Its production at different time points were further examined. Although its induction after HCoV-229E infection was observable in both NHBE and SAEC ALI, the level of IL-6 in both the apical and basal compartment in NHBE ALI was significantly lower when compared with SAEC ALI at all time points and all MOIs (Fig. 3C). Interestingly, in HCoV-OC43 infected cultures, a significantly higher level of IL-6 was also detected in SAEC ALI compared with NHBE ALI (Fig. 3D). Taken together, these data suggest the preferential induction of a more “inflamed” microenvironment in the lower respiratory tract might be a common feature of coronaviruses. It will be of interest to investigate whether SARS-CoV-2 infection induces similar immune signatures in NHBE and SAEC ALI cultures in the future.

**Figure 3.**
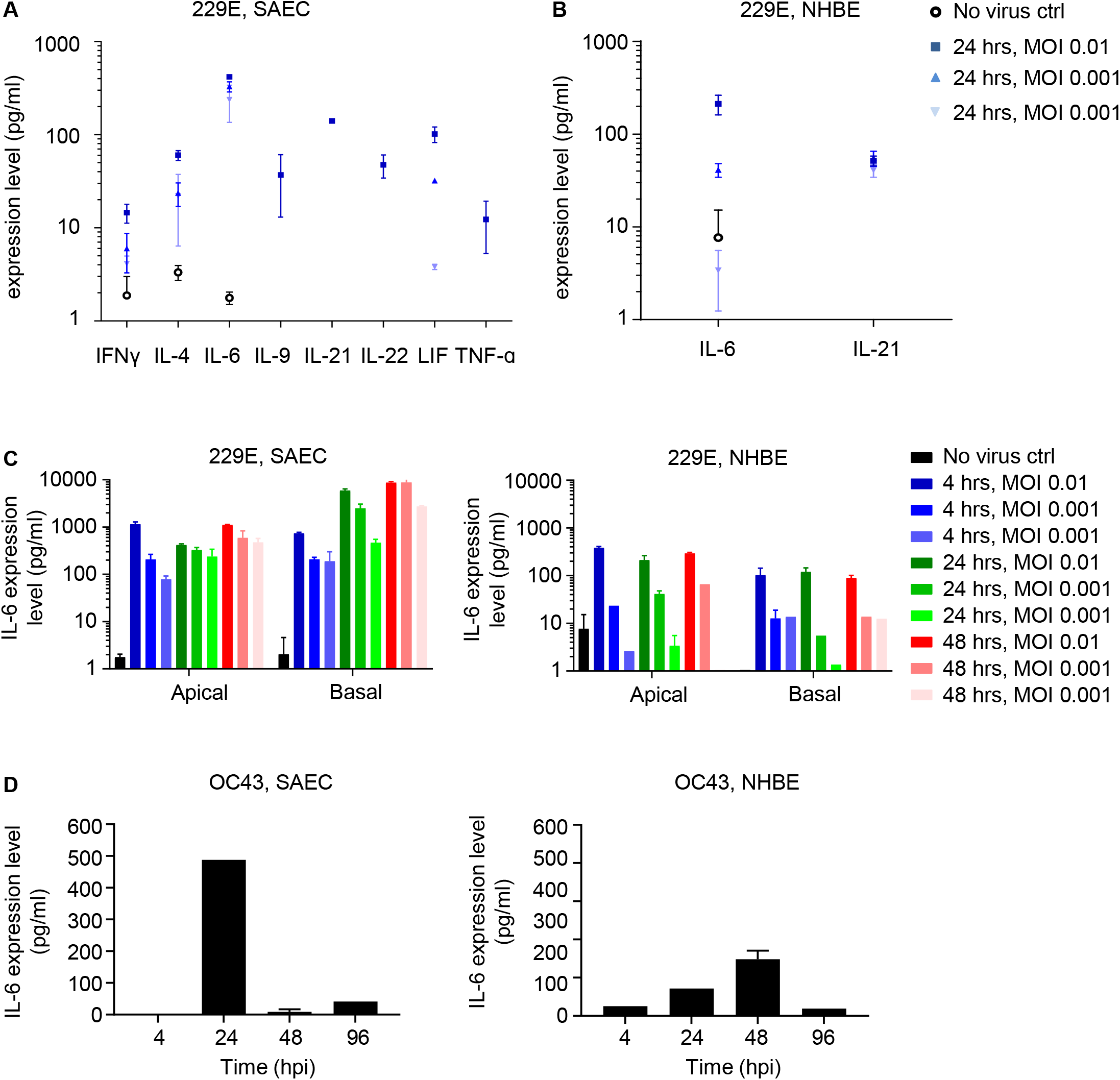
Coronavirus infection elicits differential cytokine profile in upper and lower respiratory airway. **A-B**, Apical washes from 229E infected SAEC (**A**) or NHBE (**B**) ALI culture were analyzed for the cytokine production. Infection conditions: no virus (black circle); 229E with MOI 0.01, 24 hpi (blue square); 229E with MOI 0.001, 24 hpi (blue triangle); 229E with MOI 0.0001, 24 hpi (blue inverted triangle). **C**, Apical washes or basal medium from 229E infected NHBE or SAEC ALI culture were analyzed for the IL-6 production. Infection conditions: no virus control (black); 229E with MOI 0.01, 4 hpi (dark blue); 229E with MOI 0.001, 4 hpi (blue); 229E with MOI 0.0001, 4 hpi (light blue); 229E with MOI 0.01, 24 hpi (dark green); 229E with MOI 0.001, 24 hpi (green); 229E with MOI 0.0001, 24 hpi (light green); 229E with MOI 0.01, 48 hpi (dark red); 229E with MOI 0.001, 48 hpi (red); 229E with MOI 0.0001, 48 hpi (light red). **D**, Apical washes from OC43 (MOI 0.1) infected NHBE or SAEC ALI culture were analyzed for the IL-6 production at indicated time points. All data are presented as mean ± SD of 3 independent experiments.

### HCoV-229E triggered differential transcriptional profile changes in upper and lower respiratory airways

In order to understand if the coronavirus infection caused differential transcriptional profile changes in upper and lower respiratory tracts, RNA sequencing was performed with HCoV-229E -infected NHBE and SAEC ALI. ANPEP, the entry receptor of HCoV-229E was found highly expressed in uninfected SAEC ALI but its expression level decreased dramatically after HCoV-229E infection in a time-dependent manner (Fig. 4A). This phenomenon is similar to ACE2 downregulation after SARS-CoV-2 infection (Chung et al., 2020). ANPEP expression level was much lower in uninfected NHBE ALI and did not change after HCoV-229E infection. Despite of the low ANPEP expression level, the viral yield in NHBE ALI was comparable with SAEC ALI (Fig. 2F), suggesting either a minimal level of ANPEP was sufficient for HCoV-229E entry in both models or there is an alternative entry receptor expressed on NHBE ALI to facilitate the virus entry. No statistically significant enrichment of IFN-inducible genes or apoptosis-related pathways was observed between uninfected NHBE and SAEC ALI (Fig. S2C), , indicating that in either uninfected NHBE or SAEC ALI, the interferon signaling and apoptosis were not activated. Therefore, the different histopathological changes of upper and lower respiratory airway after HCoV-229E infection are likely due to the differential host responses resulted from the viral infection. SARS-CoV-2 infection may adopt similar mechanism, considering it showed similar pathology pattern with HCoV-229E (Fig. 2A, 2C-E).

**Figure 4.**
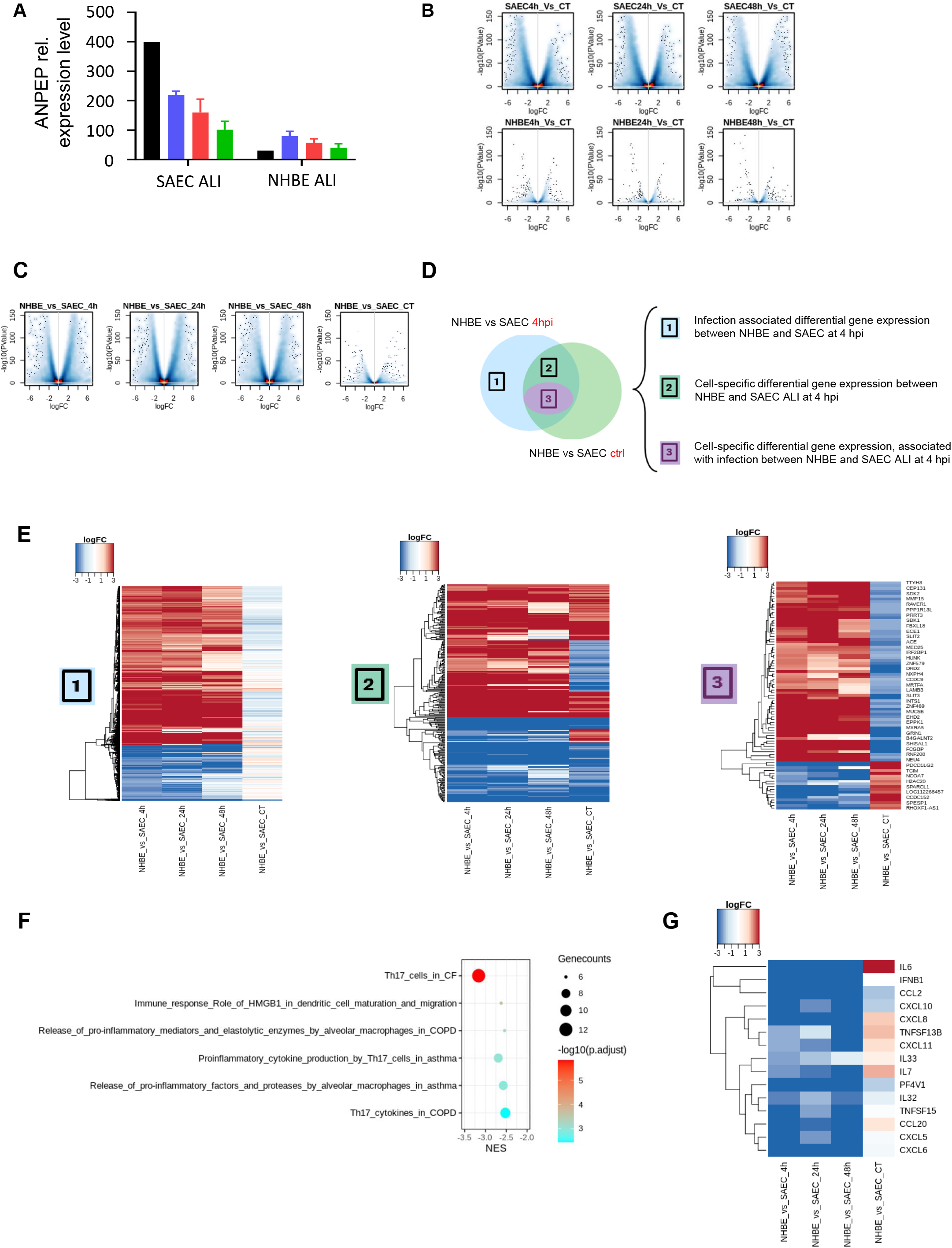
Coronavirus induces differential early immune responses in upper and lower respiratory airway. **A**, ANPEP relative gene expression level in SAEC ALI and NHBE ALI with/without 229E infection. No virus control (black bar); 229E, 4 hpi (blue bar); 229E, 42 hpi (red bar); 229E, 48 hpi (green bar). ANPEP gene counts were normalized to GAPDH. **B,** Volcano plots show differential gene expressions of SAEC infection models compared to uninfected controls at 4h, 24h and 48h timepoints (top panel), while the bottom panel represents the differential gene expression of infected NHBE models compared to uninfected controls at 4h, 24h and 48h. The x-axis is Log2 fold-change of each gene. The y-axis shows the −log10(p-value). **C,** Volcano plots depict the differential gene expression in NHBE versus SAEC models before (CT) and after virus infection (4h, 24h and 48h). **D,** Venn diagrams illustrate the overlapping genes before and after virus infection that regulated in NHBE compared to SAEC models. **E,** Heatmaps describe the log(fold-changes) of genes derived from three groups marked in Fig 4D. Red and blue indicate high and low fold-changes respectively. **F,** Dot plot describes enriched pathways of the 2719 strongly regulated genes in NHBE models compared to SAEC at 4h infection. These genes were derived from group 1 and 3 that marked in Fig 4D. The x-axis is the normalized enrichment score for each gene set and the y-axis is – log10(adjusted pvalue). Gene counts are represented by dot size. **G**, Heatmaps show the log(fold-changes) of strongly regulated human cytokine genes in NHBE models compared to SAEC before and after virus infection. Red and blue indicate high and low fold-changes respectively.

To further characterize the different host responses induced by HCoV-229E in the upper and lower respiratory tracts, we performed in-depth transcriptome analysis of NHBE and SAEC ALI before and after the virus infection. As expected, compared with the uninfected controls, the infected SAEC ALI showed a significant amount of differential gene expression at all indicated time points, whereas differential gene expression in infected NHBE ALI was sparse (Fig. 4B). Also, the abnormalities and differentiation in the regulation of gene expression in NHBE ALI and SAEC ALI occurred as early as 4 hpi (Fig. 4C). These analyses confirm that HCoV-229E infection triggered more aggressive host response in SAEC ALI at very early time of infection. We next classified these genes with differential expression between NHBE ALI and SAEC ALI at 4 hpi into three groups (Fig. 4D) and profiled the log transformed expression fold-changes of these groups using heatmaps (Fig. 4E). Differential gene expression upregulation was more frequently observed in NHBE ALI compared with SAEC ALI after virus infection (Fig. 4E). Additionally, the SAEC differential gene expression maintained the same up- or down-regulated trends as in 4, 24 and 48 hpi (Fig. 4E), suggesting that the distinct differential gene expression associated with viral infection in upper and lower respiratory airway is probably very early induced, and may persist during the infection process.

Furthermore, we conducted pathway enrichment analysis to reveal the underlying biological functions of the differential expressed genes from group 1 and group 3 (Fig. 4F). Intriguingly, majority of the top ranked pathways were directly involved in the releasing or production of cytokines and pro-inflammatory factors in the Th17 cells and macrophages. We also checked the intracellular RNA expression level of cytokine/chemokine that were shown to be up-regulated in the apical side of SAEC ALI after virus infection (Table 1, Fig. 4G), The fold change in intracellular RNA level was consistent with the amount secreted into the apical compartment and may further explain the reason why a more “inflamed” microenvironment exists in the lower respiratory tract. Apart from IL-6, To sum up, core gene sets and functional pathways that may contribute to the early immune responses during the infection of HCoV-229E was similar as reported with SARS-CoV-2 infection.

### SAEC ALI represents a physiologically relevant model for the evaluation of anti-coronavirus inhibitors

3D airway epithelial ALI model is well acknowledged as the most physiologically in vitro model for the evaluation of direct antiviral reagents (Boda et al., 2018). NHBE ALI has been employed to evaluate the anti-SARS-CoV-2 activity of RNA-dependent RNA polymerase inhibitors Remdesivir (Pizzorno et al., 2020), EIDD-1931 (Sheahan et al., 2020) and the main protease inhibitors (Fu et al., 2020; Vuong et al., 2020). However, their efficacy in SAEC ALI was not reported yet. Because SARS-CoV-2 can infect lower respiratory tract and trigger extensive epithelium damage and inflammation in lower respiratory airway, causing life-threatening pneumonia in 14% of case (Grasselli et al., 2020; Huang et al., 2020; Wang et al., 2020; Xu et al., 2020a), understanding of the efficacy of the anti-coronavirus drugs in coronavirus-infected SAEC ALI is crucial.

The anti-HCoV-229E efficacy of the main protease inhibitor GC376 was investigated in the SAEC ALI model. SAEC ALI was inoculated with HCoV-229E at an MOI of 0.01, followed by the addition of GC376 to the basal compartment which mimics intravenous drug administration. Viral replication was monitored through repeated sampling and TCID50 titration on MRC-5 cells. Treatment with GC376 at the concentration of 4 μM fully blocked the production of infectious viruses (Fig. 5A) and protected the cells from HCoV-229E virus-caused cell death, as demonstrated by the H&E staining of the intact epithelium at 72 hpi and cilia movement observation (Fig. 5B and Fig. S1C). Partial inhibition of the infectious virus production and the epithelium damage was observed with the treatment of GC376 at 2 μM. To further evaluate the potential of the combination treatment of coronavirus main protease inhibitor with RNA polymerase inhibitor, SAEC ALI cultures infected with HCoV-229E were treated with GC376 and Remdesivir simultaneously in the basal compartment. Although Remdesivir was reported to have only modest protective effect from CPEs at 0.1 μM in SAEC ALI (Zhu et al., 2021), combo of Remdesivir and GC376 resulted in sterilizing additive effect (Fig. 5C-D). All these data validate SAEC ALI model for the investigation of coronavirus infection and discovery of anti-coronavirus drugs.

**Figure 5.**
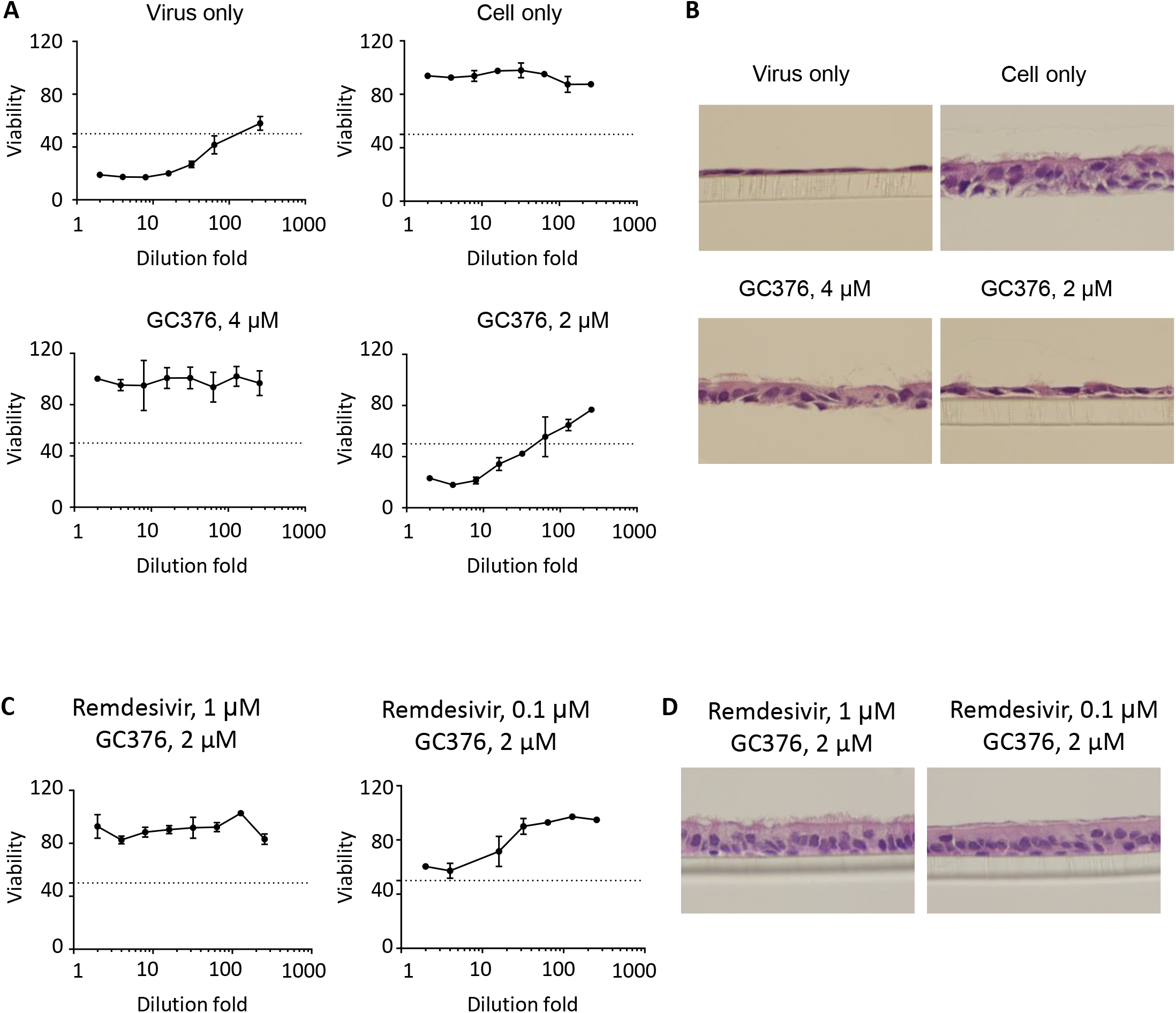
Reference antivirals inhibit coronavirus replication and protect the epithelium integrity in 3D airway epithelial ALI model. **A**, SAEC ALI cultures were infected with 229E at MOI 0.0001 followed by the treatment of GC376. Apical viral production was assessed in washes of the apical pole at Day 2. MRC5 cells were incubated with serial dilutions of the collected sample for the determination of viral titers. Dotted line indicates the TCID50/mL. **B**, H&E staining of the Fig. 6A. **C**, SAEC ALI cultures were infected with 229E at MOI 0.0001 followed by the treatment of combo of GC376 and Remdesivir. Apical viral production was assessed in washes of the apical pole at Day 2. MRC5 cells were incubated with serial dilutions of the collected sample for the determination of viral titers. **D**, H&E staining of the ALI cultures from Fig 6C. Data are representative of 3 independent experiments.

## Discussion

SARS-CoV-2-induced upper respiratory infection appears to cause mild or moderate symptoms, similar to conventional human coronaviruses that are a major cause of common colds. However, in the lower respiratory tract, COVID-19-related severe pneumonia with acute respiratory distress syndrome (ARDS) and cytokine storm syndrome challenged elderly individuals and those with comorbidities (Heymann and Shindo, 2020). The putative intrinsic differences between upper and lower airway epithelia during coronavirus infection may count for the phenotypic and clinical distinction in patients. Regarding the low physiological and functional relevance of monolayer epithelial culture to human airway during virus infection, increasing number of airway infection studies have applied the ALI models (Jazaeri Farsani et al., 2015; Stelzig et al., 2020; Wu et al., 2016). Besides the frequently used proximal airway ALIs, we at the first time reconstituted the human small airway ALI and characterized the pathophysiology and innate immune features induced by the coronaviruses with the ultimate objective to provide insight in terms of putative prognostic biomarkers and/or patient management. Our data indicate that in vitro NHBE or SAEC ALI models maintain high physiological relevance with the in vivo human respiratory airway tissues from proximal or distal part respectively. Also, the coronavirus induced immune response observed in these upper and lower respiratory airway models are similar to what has been reported in COVID-19 patients (Catanzaro et al., 2020).

Hyperactive immune responses to the SARS-CoV-2 virus infection, with the outcome of excessive release of the pro-inflammatory cytokines, was shown to be correlated with the high mortality and unfavorable prognosis of the COVID-19 patients (Ragab et al., 2020). In this study, we report the first time that both SARS-CoV-2 and HCoV-229E elicited pathophysiological and immune feature differences in human upper and lower respiratory airway. It’s been reported that in patients with COVID-19, there is upregulation of pro-inflammatory cytokines in the blood, including IL-6, TNF, and IFN-γ (Gong et al., 2020; Huang et al., 2020) and anti-IL6 antibody Tocilizumab treatment in COVID-19 showed clinical benefits (Conti et al., 2020). In accordance with these findings, we identified IL-6 associated pro-inflammatory pathways and expression level of IL-6 were significantly higher in virus-infected lower respiratory tract. Our findings that both SARS-CoV-2 and HCoV-229E could induce epithelia damage in the lower respiratory airway (Fig. 2D), is consistent with the reported lung injury or ARDS associated with severe COVID-19 patients (Chen et al., 2020). and excessive production of pro-inflammatory cytokines is considered to be one of the major contributing factors (Del Valle et al., 2020).

Novel antiviral test in the high physiologically relevant preclinical model could speed up the whole drug development progress for the COVID-19. Our data support the SAEC ALI as a promising in vitro model for the coronaviruses drug evaluation as it showed high physiology and immune feature similarity with severe COVID-19 cases. Apart from coronaviruses, we also investigated other type of respiratory viruses’ infection in our model, including respiratory syncytial virus, human parainfluenza viruses, rhinovirus and human metapneumovirus (data not shown). None of them showed similar pathophysiology effects of coronavirus, further indicating that this NHBE/SAEC ALI model could be of importance for coronavirus pathogenesis investigation.

Our data showed validated anti-viral efficacy with references compounds in SAEC ALI model and suggested the combination possibility of the polymerase inhibitor and the 3C-like protease inhibitor for the COVID-19 patients. Further investigation on the drug permeability, epithelia immune cells co-culture of SAEC ALI can be envisioned for deeper understanding of the coronavirus. In summary, a better understanding of the molecular mechanisms contributing to the coronavirus pathological differences may open new approaches for COVID-19 treatment.

## Methods

### NHBE ALI and SAEC ALI culturing

Human airway epithelial cells including bronchial epithelia cells and small airway epithelial cells were purchased from Lonza (#cc-2540s; #cc-2547s), ATCC (#PCS-300-010) or ScienCell (#7475). Cryopreserved cells were first expanded with PneumaCult™-Ex Plus Medium (STEMCELL) followed by differentiated with PneumaCult™-ALI or PneumaCult™-ALI-S Medium (STEMCELL), in 24-well plates with 6.5-mm Transwell® inserts (#38024, STEMCELL) or HTS Transwell®-96 (#100-0419, STEMCELL). The ALI models were incubated at 37°C with full medium changes in the basal chambers every 2 days until fully-differentiation is reached. The ALI models are stable for months and were usually used for virus infection and drug discovery at week 6-8 to enable the reproducible data.

### MucilAir™ and SmallAir™ culturing

AMucilAir™ and SmallAir™ were provided by Epithelix SARL (Geneva, Switzerland) and cultured at the air–liquid interface (ALI) in MucilAir™ culture medium (EP04MM) or SmallAir™ culture medium (EP65SA) (Epithelix Sàrl, Geneva, Switzerland), in 24-well plates with 6.5-mm Transwell® inserts (cat #3470, Corning Incorporated, Tweksbury, USA).

### Cell lines

The human lung fibroblast MRC5 was purchased from ATCC (USA). Cells were cultured in Eagle’s Minimum Essential Medium (Cat. 30-2003, ATCC) supplemented with 10% heat-inactivated fetal calf serum (FCS, Hyclone, USA) and 1% pen/strep (Invitrogen) at 37°C in a humidified 5% CO_2_ incubator (Thermo Fisher Scientific).

### Viruses and inoculation

Human coronavirus 229E (VR-740™) was purchased from ATCC. Coronavirus OC43 was provided by Epithelix Sàrl. Coronavirus SARS-COV-2 (French circulating strain) was provided by Epithelix Sàrl and the study was performed in VirNext (BSL3 facilities). Prior to infection, the apical side of the ALI, MucilAir™ or SmallAir™ cultures were washed once with PBS (with Ca2+/Mg2+) and transferred to a new plate containing culture medium. 100μL of 229E inoculum (MOI = 0.01, 0.001 or 0.0001), SARS-COV-2 inoculum (MOI = 0.1) or OC43 inoculum (MOI = 0.1) was applied to the apical side of the tissues for 1h at 37°C. Non-infected controls were exposed to 100 μl culture medium on the apical side for 1 h. Viruses were eliminated after the incubation period by two rapid PBS washing steps.

### Virus genome copy number

Apical washes with 300 μl of culture medium were performed for 10 min at indicated days. RNA extracted from the apical washes with the QIAamp® Viral RNA kit (Qiagen) was quantified with SuperScript™ III Platinum™ One-Step qRT-PCR Kit (Invitrogen) by quantitative RT-PCR (qPCR) in Roche Light Cycler 480 (Roche). Primers and probes used for 229E: forward primers 5′ TGG CAC AGG ACC CCA TAA AG 3′; reverse primers 5′ CAA CCC AGA CGA CAC CTT CA 3′; Probe FAM 5’- TGCAAAATTTAGAGAGCGTG-3’ BHQ. (Patent CN103993101A). Primers and probes used for SARS-Cov-2: forward primer: 5’-TGGGGYTTTACRGGTAACCT-3’; reverse primer: 5’-AACRCGCTTAACAAAGCACTC-3’; probe: 5’-FAM-TAGTTGTGATGCWATCATGACTAG-TAMRA-3’. Primers and probes used for OC43: forward primer: 5’-STC GAT CGG GAC CCA AGT AG-3’; reverse primer: 5’- CCT TCC TGA GCC TTC AAT ATA GTA ACC-3’; Probe: 5’-FAM- AGG CTA TTC CGA CTA GGT TTC CGC CTG-TAMRA-3’.

### Cell culture-based titration

Viral titers of the 229E samples were determined as log_10_TCID_50_/ml. Confluent monolayers of MRC5 cells were inoculated in duplicates with 10-fold serial dilutions of the apical wash samples and were placed at 34 °C for 3 days. Cell viability was determined using a Cell Counting Kit-8 (CCK8) kit (Dojindo, Cat.CK04) and an Envision reader (PE) according to the manufacturer’s protocol.

### Trans-epithelial electrical resistance (TEER)

300 μl of culture medium was added to the apical compartment of the tissue cultures, resistance was measured across cultures with an EVOM2 volt-ohm-meter (World Precision Instruments, Sarasota, US). Resistance values (Ω) were converted to TEER (Ω*cm2) by using the following formula: TEER (Ω*cm2) = (resistance value (Ω) − 100(Ω)) x 0.33 (cm^2^), where 100 Ω is the resistance of the membrane and 0.33 cm^2^ is the total surface of the epithelium of 24-well plate.

### H&E stain

ALI tissues were fixed with 4% Paraformaldehyde Fix Solution (#AR1068, Boster) at indicated time points overnight, followed by embedded to paraffin blocks. The paraffin-embedded tissue blocks were sectioned at 5 μm thickness on a microtome and stained with hematoxylin and eosin.

### Cytokine profiling

Apical wash and basal medium from infected ALI samples at indicated time points were measured for the cytokine production by Immune Monitoring 65-Plex Human ProcartaPlex™ Panel (#EPX650-10065-901) with The Luminex® 100/200™ System.

### RNA sample preparation and sequencing

Total RNA from ALI, MucilAir™ or SmallAir™ were extracted using Trizol LS (Thermo Fisher Scientific) according to the manufacturer’s instructions. 1 μg total RNA with RIN value above 6.5 was used for library preparation. Libraries with different indices were multiplexed and loaded on an Illumina HiSeq instrument according to manufacturer’s instructions (Illumina, San Diego, CA, USA). The sequences were processed and analyzed by GENEWIZ.

### RNAseq analysis

Paired-end RNASeq reads were mapped to the human genome (hg38) with STAR aligner version 2.5.2a using default mapping parameters (Dobin et al., 2013). The unmapped reads were later aligned to HCoV-229E genome using bowtie2 (v2.3.5) in sensitive mode (Langmead and Salzberg, 2012). Numbers of mapped reads for all RefSeq transcript variants of a gene (counts) were combined into a single value by featureCounts software (Dobin et al., 2013; Liao et al., 2014) and normalized as tpm (transcripts per million). Differentially expressed genes were identified with edgeR (v3.32.1) at a fold-change >2 and aveExpr >2.(Robinson et al., 2010)

### Gene set enrichment analysis

Gene Set Enrichment Analysis (GSEA) was performed using R package fgsea (v1.16.0) (Korotkevich et al., 2021). Gene list was ranked by logFC (NHBE compared to SAEC) in descending order. Normalized enrichment score (NES) was calculated with the internalized Molecular Signatures Database (MSigDB), MetaBase, and ImmuneSpace signatures.

### BioQC analysis

Quality control based on the enrichment of lung tissue gene signatures was performed with BioQC (v1.19.3) (Zhang et al., 2017). The BioQC scores used for visualization are log10-transformed one-sided p-values of Wilcoxon-Mann-Whitney test. The clustering and annotation of lung tissue cell types were obtained from published articles (Deprez et al., 2020 and Travaglini et al., 2020).

### Statistical Analysis

Data are presented as mean ± standard error of the mean. For statistical comparison one-way or two-way analysis of variance were performed with multiple comparison tests using GraphPad Prims software (*p < 0.05, **p < 0.01, ***p < 0.001, ****p < 0.0001).

## Author contributions

Y.Z conceived the study, performed the experiments, analysed the data, and wrote the manuscript. D.M and D.J.Z analysed and interpreted the RNA sequencing data. X.L. calculated the ANPEP expression level. Q.Z. performed H&E staining experiments. L.W. and L.G supervised the study, and wrote the manuscript.

## Financial support

This work was supported by F. Hoffman-La Roche Ltd.

## Declaration of Competing Interest

The authors are employees of F. Hoffman-La Roche Ltd.

## Notes

### Competing Interest Statement

The authors have declared no competing interest.

