## Supplemental Figure 1-3 for "A novel, anatomy-similar in vitro model of 3D airway epithelial for anti-coronavirus drug discovery"

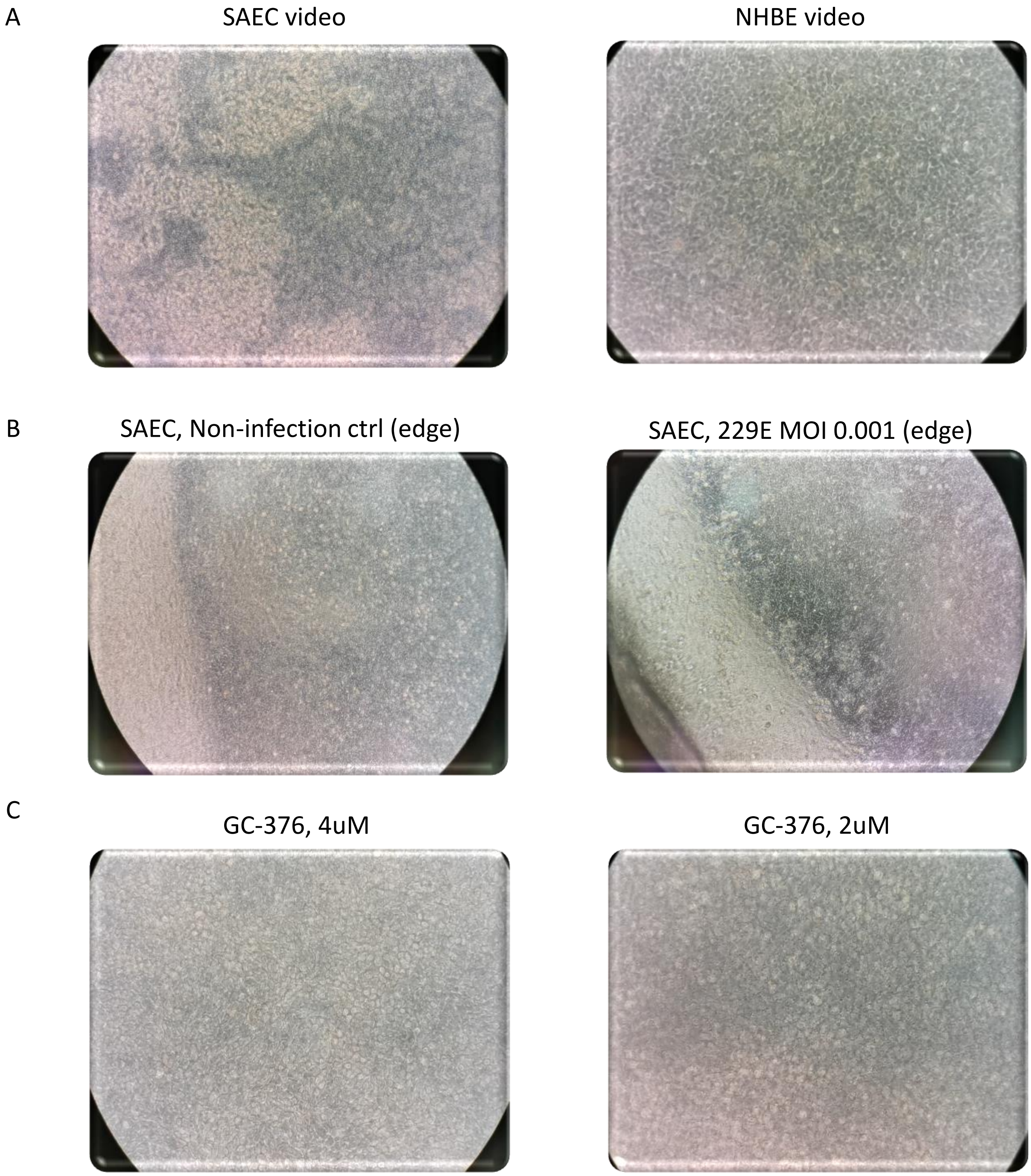


**Fig S1**, Cilia beating videos of SAEC or NHBE ALI under indicated condition.

A

B

C

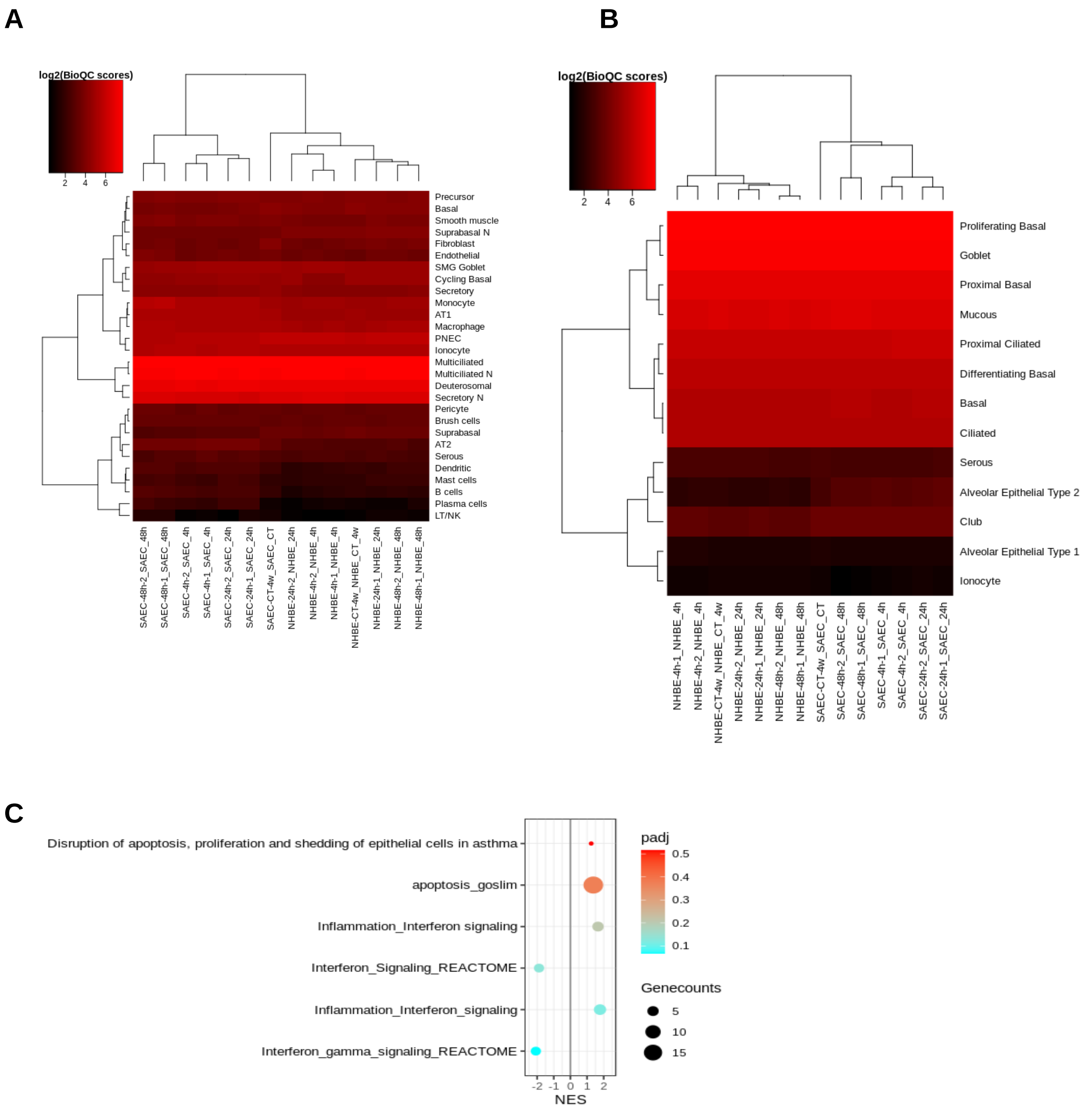


**Fig S2A-S2B**, Heatmaps depict the bioQC scores derived from SAEC and NHBE ALI models for testing the enrichment of signatures in different subtypes of lung cells (S2A--Deprez et al., 2020; S2B--Travaglini et al., 2020) . Red and black indicate high and low scores respectively. **Fig S2C**, Dot plot shows no IFN signaling or apoptosis pathway were significantly enriched in uninfected SAEC and NHBE ALI models. The x-axis is the normalized enrichment score for each gene set and the y-axis is –log10(adjusted pvalue). Gene counts are represented by dot size.

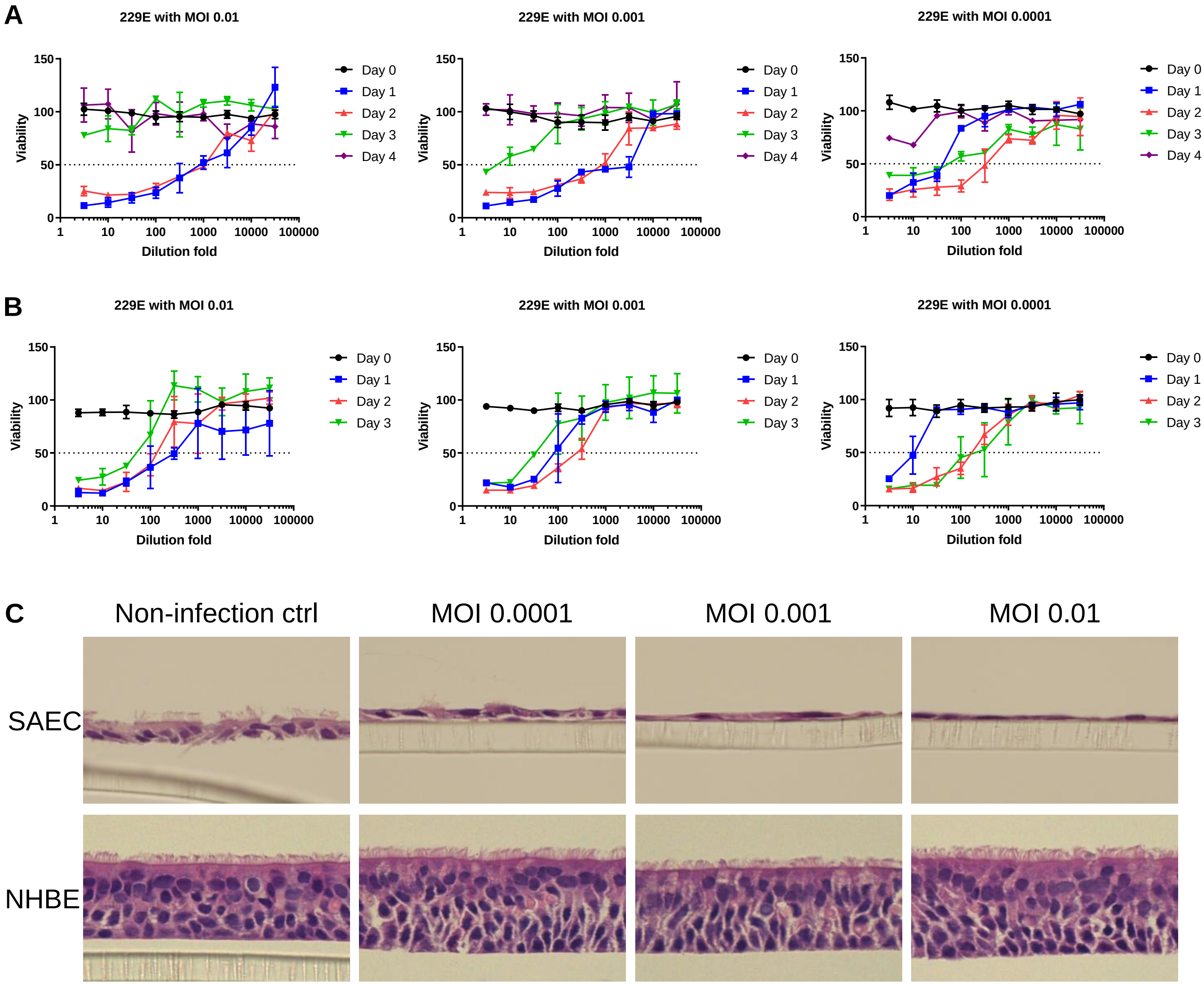


**Fig S3A-S3B**, 229E replication kinetics measured by TCID50 in MRC-5 cells with different MOIs at indicated time points in SAEC ALI (**A**) and NHBE ALI (**B**) culture. **Fig S3C**, H&E staining of the 3D cell culture at the end of infection of 229E with different MOIs in SAEC and NHBE ALI cultures.

Supplementary Figure 3
